# Spatially explicit approach to population abundance estimation in field surveys

**DOI:** 10.1101/131037

**Authors:** Nao Takashina, Buntarou Kusumoto, Maria Beger, Suren Rathnayake, Hugh P. Possingham

## Abstract

The abundance of species is a fundamental consideration in ecology and conservation biology. Although broad models have been proposed to estimate the population abundance using existing data, available data is often limited. With no information available, a population estimation will rely on time consuming field surveys. Typically, time is a critical constraint in conservation and often management decisions must be made quickly under the data limited situation. Depending on time and budgetary constraints, the required accuracy of field survey changes significantly. Hence, it is desirable to set up an effective survey design to minimize time and effort of sampling given required accuracy. We examine a spatially-explicit approach to population estimation using spatial point processes, enabling us to explicitly and consistently discuss various sampling designs. We find that the accuracy of abundance estimation varies with both ecological factors and survey design. Although the spatial scale of sampling does not affect estimation accuracy when the underlying individual distribution is random, it decreases with the sampled unit size if individuals tend to form clusters. These results are derived analytically and checked numerically. Obtained insights provide a benchmark to predict the quality of population estimation, and improve survey designs for ecological studies and conservation.

## Introduction

Estimating the abundance of populations is important for ecological studies and conservation biology [1–7], as is the role of ecosystem monitoring to observe changes in ecosystems [8–10]. In conservation, such knowledge helps one to estimate the risk of extinction of species [11, 12], and to implement effective conservation actions [13].

While methods for statistically inferring population abundance with existing spatial data are well developed [4–6, 14, 15], information on the abundance of threatened or rare species is often rather limited and biased given budgetary constraints and access to remote sites [16, 17]. For example, Reddy and Davalos [16] examined an extensive data set of 1068 passerine birds in sub-Saharan Africa, and they found that data on even well-known taxa are significantly biased to areas near cities and along rivers. Typically, time is one of the critical constraints in conservation areas facing ongoing habitat loss and environmental degradations [18]. In such cases, management decisions must be made quickly under the data limited situation with often limited knowledge of a system [13, 19, 20]. On the other hand, for many ecosystem monitoring programs, corrected data must be accurate enough to be able to detect ecological change [9]. Hence, given time and budgetary constraints, it is desirable to set up an effective survey design to minimize time and effort of sampling to satisfy required accuracy.

Ultimately, we face a trade-off between accuracy, time, and money. To tackle this trade-off, and provide advice to people designing a spatial sampling approach, we need a theory that can handle different sampling methods, choice of sampling unit size, and fraction of sampled region. Most previous approaches are spatially implicit e.g., [5,6,14,15,21], and it is therefore not straightforward to compare the effect of different survey designs. For example, as in the previous studies just mentioned above, the negative binomial distribution (NBD) is the most frequently used distribution for describing the underlying biological distribution of a species. In the NBD, the parameter characterizing the degree of spatial aggregation is scale dependent, and needs to be calibrated for each sampling unit scale. However, this procedure is not intuitive, as the parameter characterizing aggregation is usually inferred by observed data rather than biological mechanisms [14].

To address this challenge, here we use a spatially explicit approach, which enables us to explicitly and consistently compare the effect of different sampling schemes across sampling unit scales. Specifically, we examine random sampling and cluster sampling [22, 23] as sampling schemes, due to their simplicity of implementation. Moreover, cluster sampling reflects existing geographically biased sampling to some extent. These sampling schemes are combined with spatial point processes (SPPs), a spatially-explicit stochastic model to reveal effects of different survey regimes as well as ecological factors on the performance of population estimate. SPPs are often applied in ecological studies due to their flexibility and availability of biological interpretations [24–28]. Many of their examples come from studies of plant communities [24, 25, 27, 28], but include also study of coral communities [29], where spatial aggregation patterns are typically observed [30, 31]. Hence, field surveys of plant and coral species are the potential applications in our framework.

Using a spatially-explicit approach, we revealed new insights into population estimation with different survey designs as well as ecological regimes. The accuracy of the abundance estimation through random sampling varies with both ecological factors, such as the spatial distribution pattern of individuals, and survey design. Although the spatial scale of sampling does not affect estimation accuracy when the underlying spatial pattern is random, it decreases with the size of sampling unit if individuals tend to form clusters. However, under both random and clustered distributions, the relative standard error of abundance estimation decreased with the number of individuals, and the fraction of a region sampled. Obtained these insights will help to improve field survey designs for both ecological studies and conservation planning.

## Methods

In this analysis, we consider a situation where there is no prior spatial data available to infer the distribution and abundance of a target species. We assume that our estimate of population size is based only on field surveys where a sampled fraction *α* (0 *< α ≤* 1) of the region of concern, *W*, is surveyed using a sampling unit size, *S* (Fig. 1). We focus on a case where no sampling error occurs in each sampling unit, suggesting that a sampling unit size should be chosen to satisfy this requirement in practice, and it may vary for sampling in different systems. For example, such an area may be larger for counting plant species compared to counting coral species due to different visibility and accessibility.

First, we introduce an estimator of population abundance, its expected value and variance, which explicitly accounts for the effect of sampling unit size. Next, we explain some basic properties of spatial point processes (SPPs),and models to describe spatial distribution patterns of individuals. Using computer-generated spatial distribution patterns, we test our analytical results formula for population estimation.

**Figure 1:**
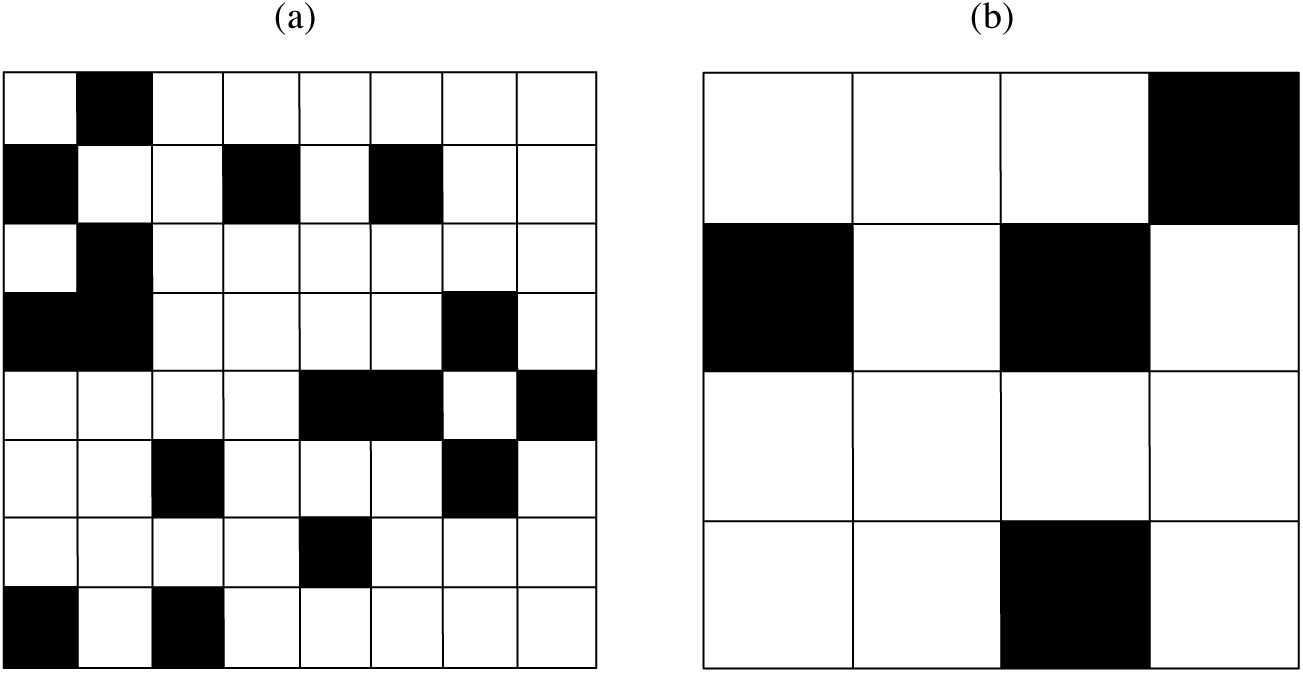
Example of simple random sampling with (a) smaller, and (b) larger sampling unit size. The whole region of concern is divided into sampling unit with equal size, and a certain fraction *α* is randomly sampled (shaded unit) without replacement, where all sampling units have the equal probability to be chosen. No or sufficiently small measurement error is accompanied by each sampling trial. Essentially, applying larger sampling unit corresponds to a cluster sampling. The examples show the case of *α* = 0.25.

### Survey design

Given parameters specifying the survey design noted above, a simple random sampling (SRS) without replacement [23] is conducted for correcting count data (Fig 1). In the SRS without replacement, all the sampling units have an equal probability to be chosen and units already sampled once are not chosen again. The number of sampling units, *N*_*t*_, and the sampled units, *N*_*s*_, change with a sampling unit size, *S*. We assume all the sampling units have an equal size. As applying larger sampling unit sizes, the degree of the geographical sampling bias increases when the fraction of a sampled region is small. Essentially, it corresponds to one-stage cluster sampling [23], where either all or none of the parcel of the area within the larger sampling unit are in the sample. It is worth noting, however, that the sampling unit size is arbitrarily chosen in ecological surveys; therefore the degree of cluster sampling is relative: any SRS can turn to be a cluster sampling if it is compared to SRS with a smaller sampling unit size. In this article, we simply use the simple random sampling and cluster sampling by implying using relatively *small* and *large* sampling units.

### Population estimator

Following the data collection, we apply the following unbiased linear estimator of the population abundance in the region of concern *W*, *n*(*W*) [22, 23],

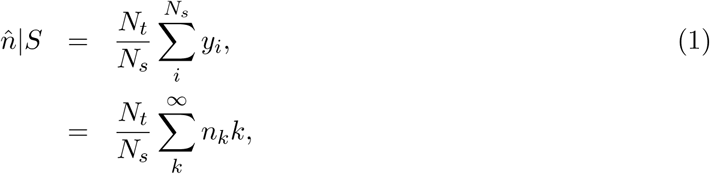

where, 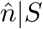 is the estimated population abundance given sampling unit size *S*, *y*_*i*_ is the number of sampled individuals at *i*th sampling trial, and *n*_*k*_ is the frequency of the sampled units holding *k* individuals. Note *y*_*i*_ and *n*_*k*_ change depending on the sampling unit size and underlying spatial point patterns. In the SRS without replacement, the frequency *n*_*k*_, given the number of sampled units *N*_*s*_, is only the probability variable, following a multivariate hypergeometric distribution *p*(*n*_*k*_*|S, N*_*s*_) with the mean *N*_*s*_*p*(*k|S*). Hence, the average population estimation 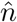 is

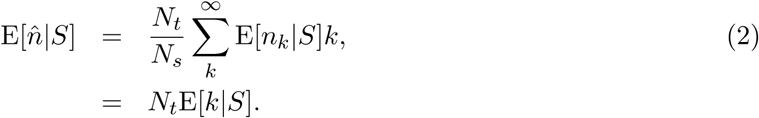

The variance of population estimate under the SRS without replacement is obtained by multiplying the finite population correction (fpc) := (*N*_*t*_ *- N*_*s*_)*/*(*N*_*t*_ - 1) [22] by the variance under the SRS with replacement:

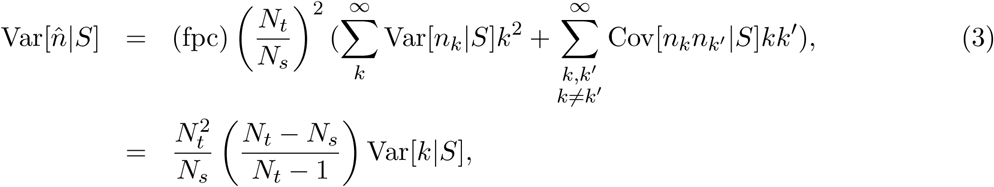

where, the fact that the probability *p*(*n*_*k*_*|S, N*_*s*_) follows a multinomial distribution with Var[*n*_*k*_*|S*] = *N*_*s*_*p*(*k|S, N*_*s*_)(1 *-p*(*k|S, N*_*s*_)) and Cov[*n*_*k*_*n*_*k*_*’ |S*] = *-N*_*s*_*p*(*k|S, N*_*s*_)*p*(*k′|S, N*_*s*_) (*k* ≠ *k′*) are used. Therefore, the variance of the abundance estimate is determined by a constant multiplied by variance of individual numbers in the sampling unit.

### Spatial distribution of individuals

To account for explicit spatial distributions of individuals, we use spatial point processes (SPPs) [24, 28]. The underlying models used in our analysis are the homogeneous Poisson process and Thomas process, generating random and cluster distribution patterns of individuals, respectively. Properties of these processes are found in the literature (e.g., [24, 28, 32]) and, hence, we only introduce the properties relevant to our questions.

### Homogeneous Poisson process

One of the simplest class of SPPs is the homogeneous Poisson process where the points (i.e. individuals) placement are randomly determined within the region of concern and the number of points given in the region *R*, *n*(*R*), is according to the Poisson distribution with an average *μ*_*R*_:

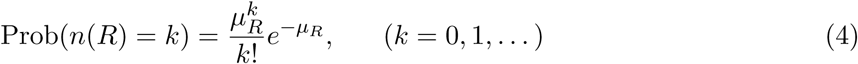

where, *μ*_*R*_ is known as the intensity measure [24, 28] defined by

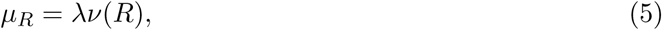

where, *λ* := *n*(*W*) */ν*(*W*) is the point intensity of the whole region concerned *W*, and *ν*(*R*) is the area of region *R*.

### Thomas process

The Thomas process, characterizing the clustering pattern of individuals, belongs to the family of Neyman-Scott processes [24, 28]. It provides a general framework to address spatial ecological patterns since almost species are clumped in nature [33]. In addition, the Thomas process is relatively amenable to an analytical approach [24, 25, 27, 28]. The Thomas process is obtained by the following three steps:

1. Parents are randomly placed according to the homogeneous Poisson process with a parent intensity *λ*_*p*_.
2. Each parent produces a random discrete number *c* of daughters, realized independently and identically.
3. Daughters are scattered around their parents independently with an isotropic bivariate Gaussian distribution with variance *σ*^2^, and all the parents are removed in the realized point pattern.

The intensity of the Thomas process is [28]

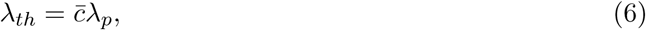

where, 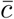 is the average number of daughters per parent. For the sake of comparison between the population estimation of the two SPPs under the same average number of individuals, we chose the intensity of the Thomas process so as to have the same average individuals within the region of concern *W*. Namely, the parameters *λ*_*p*_ and 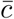 satisfy

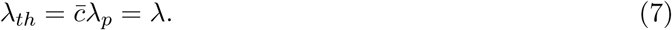

We also assume that the number of daughters per parents *c* follows the Poisson distribution with the average number 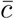

## Results

The total number of sampling units and sampled number is *N*_*t*_ = *ν*(*W*) */ν*(*S*) and *N*_*s*_ = ⌊ *αN*_*t*_ ⌋, where ⌊ *x* ⌋ is the maximum integer not larger than *x*. We are here interested in how the estimations of population total deviate from the true value, and its relative degree compared to the true value. Therefore, one of the quantities to show these effect may be

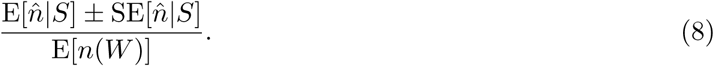

Note in the analysis below, we use ⌊ *αN*_*t*_ ⌋ = *αN*_*t*_ for simplicity, but this effect is rather small when *αN*_*t*_ is sufficiently large.

### Population estimation under the homogeneous Poisson distribution

For the homogeneous Poisson process, Var[*k|S*] is equivalent to the variance of the Poisson process with average *λν*(*S*). Therefore, by substituting this expression into Eq. (3) and with some algebra, we obtain the SE of the homogeneous Poisson process

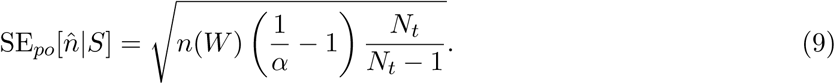

When the total number of sampling units is sufficiently large (*N*_*t*_ ≫ 1), we obtain

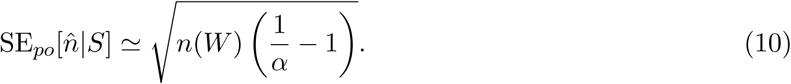

Under such circumstances, the standard error of the abundance estimation is only the function of the expected population total existing in the concerned region *n*(*W*) and the sampling fraction *α*; and does not depend on the sampling unit size. Therefore, we can write 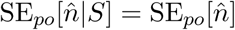. Due to the term *n*(*W*)^1*/*2^ in 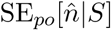, the relative variation from its average decreases with the factor (1*/α -* 1)^1*/*2^*n*(*W*)^-1*/*2^. These results are confirmed by numerical simulations, and they show good agreement with analytical results (Fig. 2, 3).

### Population estimation under the Thomas process

For the Thomas process, deriving an analytical form of the variance of individuals given a spatial scale of sampling Var[*k|S*] is challenging, although the probability generating functional of the Thomas process is known, e.g., [28]. Here we apply an approximated probability distribution of the Thomas process for the sake of obtaining an explicit form of 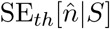. Assuming that all parents within the region *S*′, where parents can potentially supply daughters to a region *S*, given the same number of daughters, we derive an approximated 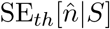 as (Appendix).

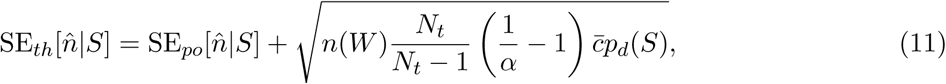

where, *p*_*d*_(*S*) is the probability that an individual daughter produced by a parent situated a surrounding region of *S*, *S*′, falls in the region *S* (Appendix). Therefore, the standard error of the Thomas process, 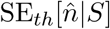, can be described by the sum of the standard error of the Homogeneous Poisson process and a term characterizing the degree of cluster of the Thomas process. The second term of Eq. (11) increases with the degree of the clustering. Because *p*_*d*_(*S*) approaches to 0 as *ν*(*S*) becomes small and approaches to 1 as *ν*(*S*) becomes large (see Eq. (A.2) and Fig. A.2 in Appendix), Eq. (11) can be described by simpler form when *p*_*d*_(*S*) *∼* 0 and *p*_*d*_(*S*) *∼* 1. Especially, if 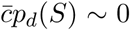 and *N*_*t*_ ≫ 1, Eq. (11) coincides with 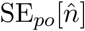, where 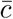Similarly, as *p*_*d*_(*S*) increases with the sampling unit size, applying a geographically biased sampling increases the standard error. It is the clear difference from the result obtained under the random spatial pattern of individuals. Biologically speaking, species with a high fecundity, 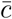, and smaller dispersal distance of daughters, *σ*, increases the 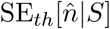, and vice versa.

**Figure 2:**
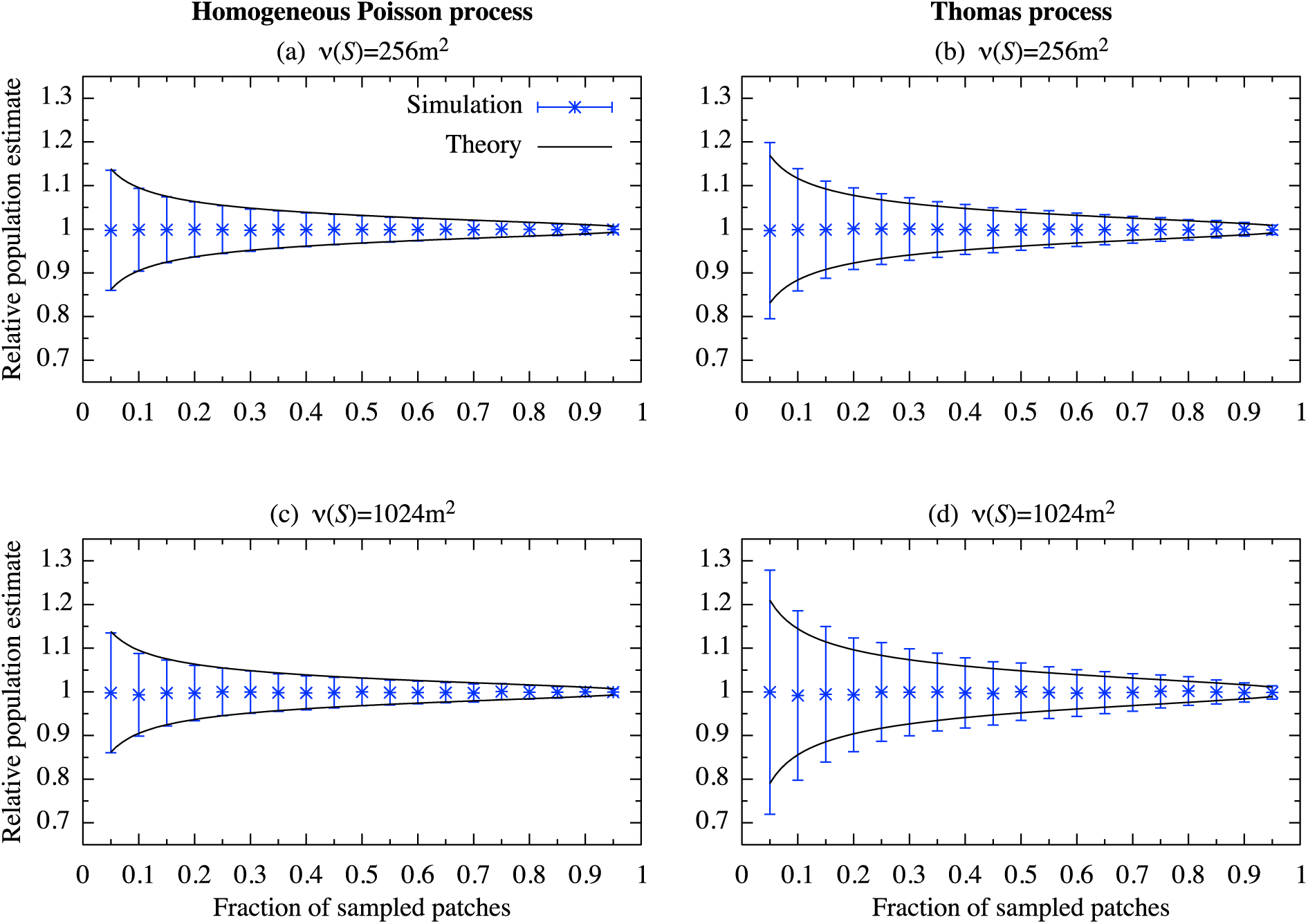
Relative value of the population estimate with the average individuals E[*n*(*W*)] = 10^3^. The parameter values used are 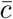, *σ* = 10, and *ν*(*W*) = 2^20^m^2^ (1024m*×*1024m).

**Figure 3:**
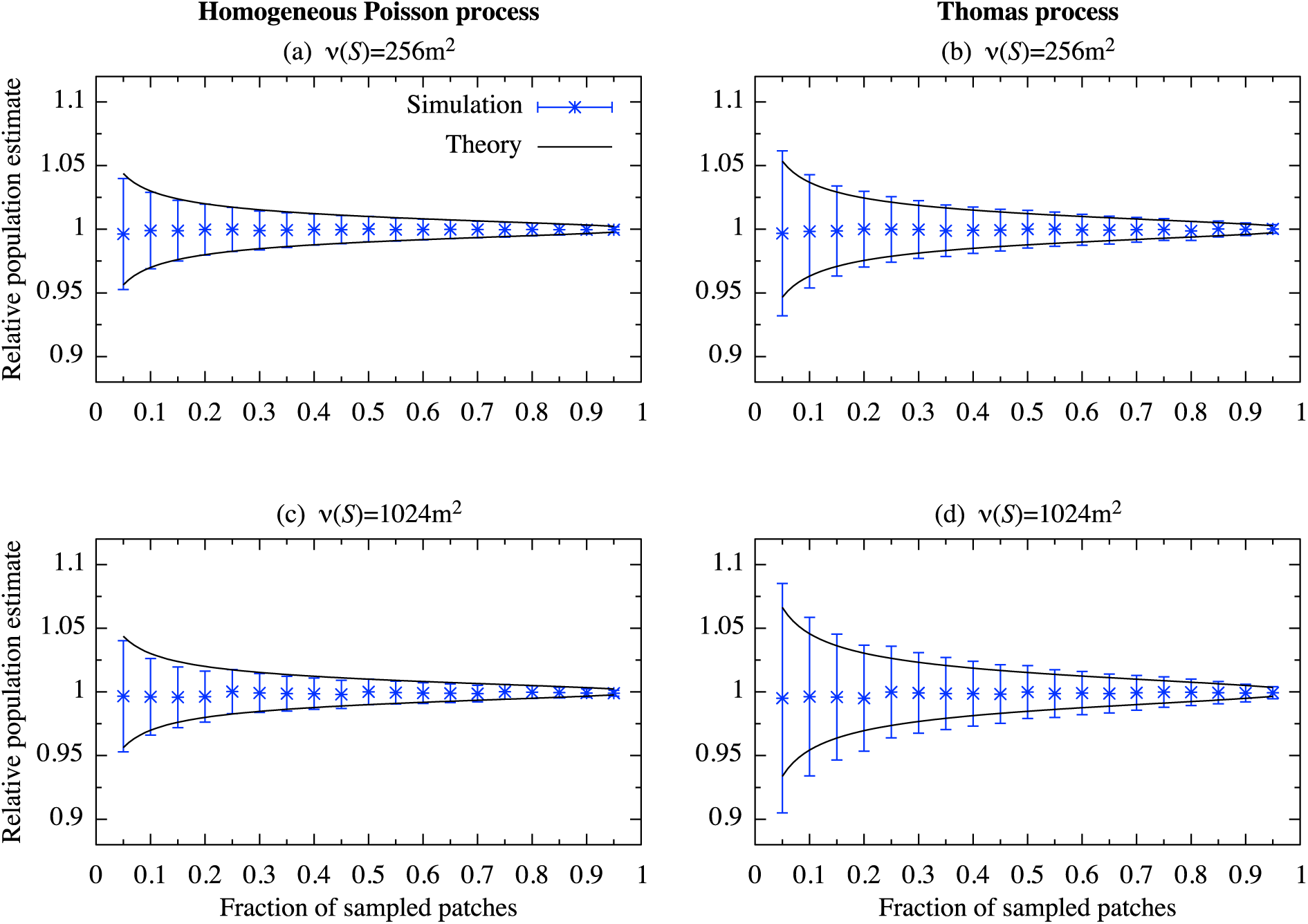
Relative value of the population estimate with the average individuals E[*n*(*W*)] = 10^4^. The parameter values used are the same as in Fig. 2.

The approximated 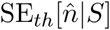, Eq. (11), shows good agreement with the values obtained by the numerical simulations, except where the fraction of the sampled patch is small (Fig. 2, 3). However, its relative deviation from the numerical values are decreased as the average number of individuals E[*n*(*W*)] increases (Fig. 3).

## Discussion

We examined a method for spatially-explicit population estimation combined with spatial point processes (SPPs) to reveal effects of different survey regimes as well as ecological factors on the performance of population estimate. By assuming the random and clustering placements of individuals as underlying ecological distribution patterns, that is, the homogeneous Poisson and Thomas process, we analytically show that the individual distributions and sampling schemes, such as random sampling and cluster sampling, change significantly the standard error of the abundance estimate. In our sampling framework, increasing the sampling unit size corresponds to an increase of geographical bias of the sampling. Typically, we find that the standard error of the abundance estimate is insensitive to the sampling unit size applied when the underlying ecological distribution is the homogeneous Poisson process. It is however individuals of most species are typically aggregated [30,34] unless the abundance of a species is low [34]. Instead, when the underlying ecological pattern is described by the Thomas process, the standard error is increased with the size of sampling unit. That is, under clustered ecological distributions, the standard error is increased as the degree of clustering sampling increases. We also show that the standard error of the population estimate increases with the parameter controlling the degree of clustering of individual distributions 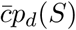(Eq. 11). In addition, although for both ecological distribution patterns our results suggest that absolute value of the standard error increases with the number of individuals, the relative standard error decreases with the factor proportional to *n*(*W*)^-1*/*2^. This means that for a species with a smaller number of individuals, the accuracy of the population estimation becomes lower. Although the homogeneous Poisson process may be a crude assumption for a distribution pattern of individuals, its sampling outcomes coincide with that of the Thomas process when the number of sampling unit is large and 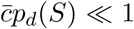. This occurs if the fecundity of a species of concern 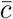 is sufficiently small, or the probability that an individual produced by a parent within a region *S*′, where a parent potentially supplies her daughter to a sampling unit *S*, is sufficiently small.

In practice, simple random sampling may not easily be conducted due to time consuming, costly, and different accessibility to site [16,23,35]. However, simple random sampling may be a reasonable option when available information on the underlying species distribution patterns is limited [23], and it may enable us to obtain more reliable data since extensive samplings in inaccessible region may also lead to new discoveries [16]. Alternatively, cluster sampling, which causes a geographical sampling bias but less expensive, is of often the favored survey design [16,23]. This may be suitable for the data collection of a species needed for quick conservation action at a cost of accuracy of data.

The central assumption in the analysis is that measurement error (i.e. imperfect detection) does not occur or is sufficiently small. Although many empirical studies have adopted this assumption [36], imperfect detection is also frequently observed even in sessile organisms like plants (e.g. [37, 38]). If searching time is fixed, rates of imperfect detection can be expected to increase with survey area [39]. This indicates that the sampling unit size should be chosen while taking the scale-dependency of the imperfect detection into account. Further studies are needed for more quantitative arguments about this issue.

Here we investigate the population estimation across sampling unit sizes under random or clustered ecological distribution patterns. Our results provide general insight into ecological survey design such as how the sampling unit size used and ecological spatial distribution patterns affect the estimation accuracy. For both ecological and conservation application in mind, our sampling framework is kept as general as possible. Therefore, it allows one to further extend our framework to handle more complex situations where, for example, the concerned region holds multiple institutions with different sampling unit sizes or a budgetary constraint is explicitly taken into consideration.

## Acknowledgements

We would like to thank T. Fung and B. Stewart-Koster for their thoughtful comments. NT and BK were funded by the Program for Advancing Strategic International Networks to Accelerate the Circulation of Talented Researchers of the Japan Society for the Promotion of Science, and they acknowledge the support for coordinating the research program from Dr Yasuhiro Kubota and Dr James D. Reimer.

## Appendix: Approximated first and second moment of the Thomas process

Here, we derive an approximated probability distribution function (pdf) given area *R* of the Thomas process. Let the whole region potentially contribute to the region *R* be *R*′ = *R* + *R*_*out*_, where *R*_*out*_ is the region where parents outside the region, *R*, can potentially supply daughters to the region *R* (Fig. A.1). Here. We assume that all parents give the same number of daughters within the small region, corresponding to an assumption that the variance in the numbers of daughter between parents is small. By the procedures to obtain the Thomas process, the probability *n* individuals found given region *R* may be described as follows:

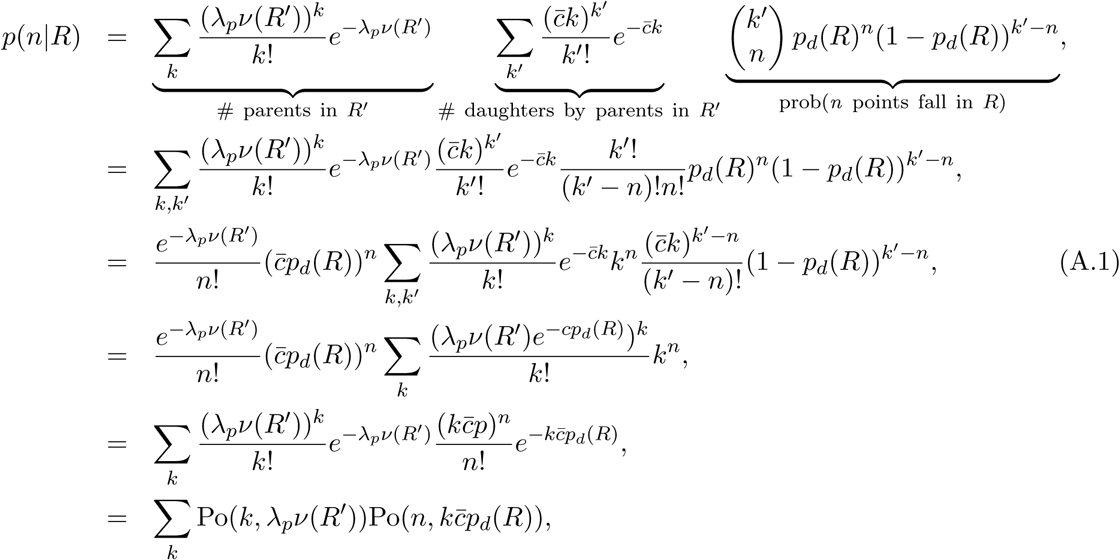

where, *p*_*d*_(*R*) is the probability that an individual daughter produced by a parent within *R*′ falls in *R*:

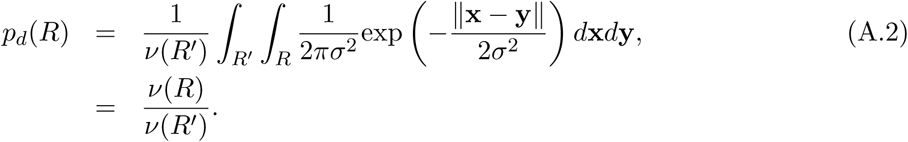

By referring Fig. A.1, *ν*(*R*′) is calculated as

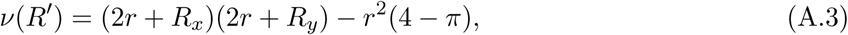

where, *r* is the distance that on average a fraction *u* of daughters scattered by the parent (placed center) are covered. *r* is calculated by converting the expression of the isotropic bivariate gaussian on cartesian coordinates, 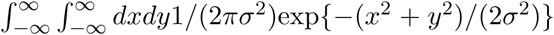, to the one on the polar coordinates, and solving about *r*

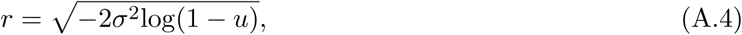

where, in the analysis, we set *u* = 0.99. An example functional form of Eq. (A.2) is shown in Fig. A.2.

**Figure A.1:**
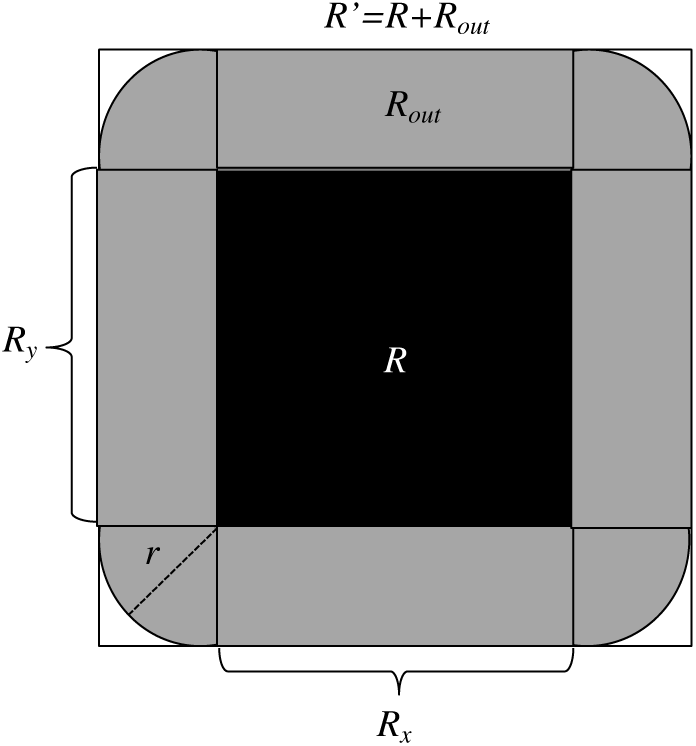
*R* is the concerned region with area *R*_*x*_ *× R*_*y*_. Parents outside *R* with a distance less than *r* from the edges of *R* (parents in *R*_*out*_) may also contribute to the number of daughters in the concerned region *R*. The whole region where parents can supply daughters to *R* is *R*′ = *R* + *R*_*out*_.

**Figure A.2:**
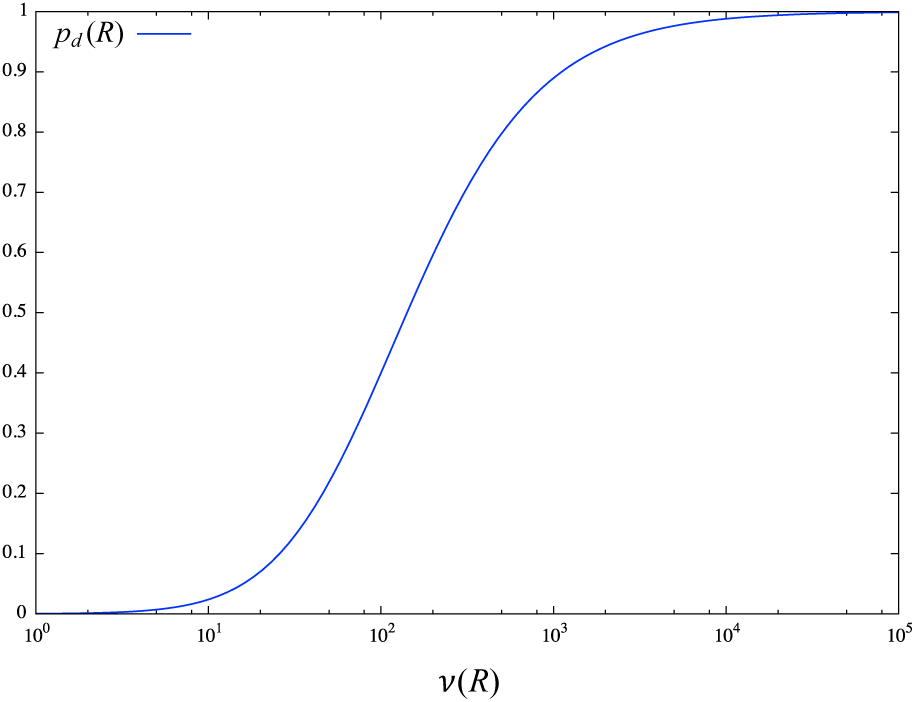
Probability that an individual daughter produced by a parent within *R*′ falls in *R*. The parameter values are the same as used in analysis. Namely, *u* = 0.99, *σ* = 10, and these give a value *r* = 30.35.

Using Eq. (A.1), we calculate the first moment and the second moment of point number *k* in region *R* as

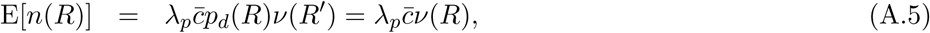

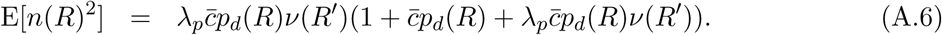

Using Eqs (3), (A.5), and (A.6) and the assumption 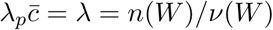, we calculate Eq. (11).

